# Enhancing the cell-free expression of native membrane proteins by in-silico optimization of the coding sequence – an experimental study of the human voltage-dependent anion channel

**DOI:** 10.1101/411694

**Authors:** Sonja Zayni, Samar Damiati, Susana Moreno-Flores, Fabian Amman, Ivo Hofacker, Eva-Kathrin Ehmoser

**Author notes:** These authors contributed equally to this work (Sonja Zayni, Samar Damiati).

## Abstract

The investigation of membrane proteins, key constituents of cells, is hampered by the difficulty and complexity of their in vitro synthesis, of unpredictable yield. Cell-free synthesis is herein employed to unravel the impact of the expression construct on gene transcription and translation, without the complex regulatory mechanisms of cellular systems. Through the systematic design of plasmids in the immediacy of the start of the target gene, it was possible to identify translation initiation and the conformation of mRNA as the main factors governing the cell-free expression efficiency of the human voltage dependent anion channel (VDAC), a relevant membrane protein in drug-based therapy. A simple translation initiation model was developed to quantitatively assess the expression potential for the designed constructs. A scoring function is proposed that quantifies the feasibility of formation of the translation initiation complex through the ribosome-mRNA hybridization energy and the accessibility of the mRNA segment binding to the ribosome. The scoring function enables to optimize plasmid sequences and semi-quantitatively predict protein expression efficiencies.

## Introduction

Understanding structure and function of membrane proteins is key in many biological processes, yet faces numerous issues. Membrane proteins are notoriously difficult to synthesize: in cells, these are usually expressed in low amounts, and their expression profile is heavily controlled as part of regulatory processes and transduction. Besides, in-cell expression of recombinant membrane proteins only works for those proteins that do not significantly alter the physiology of their hosts. The characterization of membrane proteins is no less difficult: the structural integrity of membrane proteins is hard to preserve in extracellular conditions, and function may be lost if proteins are removed from their native membranes.

The production of membrane proteins outside living cells circumvents many of the issues of in-cell synthesis. ^[1, 2]^ Cell-free synthesis uses cell lysates to in situ generate rightly folded membrane proteins ^[3, 4]^ from exogeneous mRNA or DNA, which can be directly incorporated into artificial membranes. ^[5]^

Yet cell-free and in-cell synthesis face a common challenge. In both the design of the plasmid vector is crucial. This genetic construct lodges the sequences of the transcription promotor, of the ribosomal binding site, RBS, and occasionally of translation enhancers in addition to the target gene. ^[1, 6]^ The sequence layout, particularly in the vicinity of the gene’s initiation or start codon, has become the quintessence of cell-free protein expression and yet it has not been fully exploited in optimizing constructs for protein expression. The coding region adjacent to the start codon remains untapped in both *in-silico* ^[7,8]^ and wet-bench design of constructs, and finding a working construct is to date mainly based on trial and error

Herein we present a rationalized approach to the generation of constructs for the expression of wild-type, human membrane proteins in prokaryotic cell-free systems, that includes alterations in the coding sequence proximal to the start codon. As a relevant case example, we chose the human voltage dependent anion channel or VDAC; a small, 285-amino acid-long protein (M_w_=31 kDa), that is predominantly found in the mitochondrial outer membrane ^[9,10]^. VDAC forms cylindrical channels across the membrane of diameter 20–30 Å, allowing the passage of ions and small molecules ^[11, 12, 13],^ and is involved in various pathophysiological mechanisms.

## Experimental Section

### Cloning and purification of plasmids

Cloning was performed with Gateway® recombination cloning technology (Invitrogen, Thermo Fisher Scientific, Waltham USA). Eight forward and one reverse primers were designed, see ^**[30]**^. The DNA of VDAC (855 base pairs), was amplified by PCR (Biometra Thermocycler, Analytik Jena, Jena DE), with Phusion DNA polymerase (Thermo Fisher) and vector pQE30-VDAC as template. All PCR products were purified with the MinElute PCR purification kit (Qiagen, Venlo, NL). Gateway® recombination was performed with enzyme mixes BP Clonase II and LR Clonase II according to manufacturer’s instructions (Invitrogen, Thermo Fisher Scientific). BP reactions were carried out with the purified fragments and the entry vector pDONR221. LR reactions were performed with entry clones from individual bacterial colonies and destination vectors pDEST14 and pDEST17. BP and LR products were subsequently transformed into *E. coli* strains DH5α and Top 10 (Invitrogen). Positive clones were identified by in situ PCR (RedTaq Master Mix, Sigma-Aldrich, St Louis, USA). Plasmid DNA was then purified with the QIAprep Spin Miniprep Kit (Qiagen) and examined through digestion with restriction enzymes *EcoRI*/*HindIII* and *PstI/Xhol* for DONR and DEST vector constructs, respectively (Termo Fisher). Sequencing of VDAC gene inserts for DONR and DEST vectors was performed with the VDAC specific and T7 promotor/T7 terminator primers (LGC Genomics, Berlin, DE; Microsynth, Balgach, CH), respectively. Plasmids were purified with Midi preps (Qiagen Plasmid Midi Kit or innu PREP Plasmid MIDI Direct Kit, Analytik Jena).

### Cell-free synthesis

Reactions were performed with two different kits, the S30 T7 High-yield protein expression system (Promega, Fitchburg, USA) and the PURExpress R in Vitro Protein Synthesis Kit (New England BioLabs, Ipswich, USA) according to the manufacturer’s instructions. The results herein reported refer to those obtained with the second kit, as it proved the most effective. 250 ng of plasmid, 0.2 μl Ribonuclease inhibitor (RNasin, Promega) and 0.4 μl of FluoroTect TM Green_Lys_ were added to PURExpress extracts to a reaction volume of 10 μl. After a 2 hour incubation at 37°C, 10 μl of sample dilution buffer (LDS sample buffer reducing agent, Invitrogen, Thermo Fisher) was added to the mixture. Protein denaturation in the diluted samples was conducted at 70°C for 10 min before electrophoresis.

### SDS-Page and Western Blot

The denatured samples were loaded into 10% precast gels (Invitrogen, Thermo Fisher). Electrophoresis was conducted at a constant potential of 200V for 45 minutes and imaged immediately after with a Safe Imager 2.0TM Blue Light Transilluminator. The emission fluorescence at ~470 nm of the fluorescent lysine accounted for the optical visualization of the protein bands. Thereafter Coomassie staining was performed with SimplyBlueTM SafeSTain solution (Invitrogen Thermo Fisher) on the same gels. Transfer to PVDF membranes (iBlot®, Thermo Fisher) was conducted on a second gel. Immunodetection of proteins was carried out in an InfraRed Imager (Odyssey® Infrared Imaging System, LI-COR Biosciences, Lincoln, USA), using rabbit monoclonal anti-VDAC (Cell Signaling Technology, Cambridge, UK), or anti-6x HIS-tag (Gen Tex) as primary antibody, and goat anti-rabbit IRDye 680 (LI-COR) as secondary antibody. PageRulerTM Plus Prestained Protein Ladder (Thermo Fisher) were used as standard.

### RNA detection and quantitative PCR (qPCR)

Levels of RNA were measured with a ND-10000 Spectrophotometer (Nanodrop Technologies, Wilmington USA) on RNA-isolated samples ^**[14]**.^ For qPCR, ~650 ng of isolated RNA was reversed-transcribed into cDNA with the iScript TM Select cDNA synthesis kit and random primers (Bio-Rad, Hercules, USA). qPCR was performed in a 48-well, MiniOpticon Real-Time PCR System (Bio-Rad) on sample triplicates (20 μl total reaction volume) ^**[14]**.^ SsoAdvancedTM Universal SYBR Green Supermix (Bio-Rad), was used to prepare the master mix for each primer.

### Calculation of δ and in-silico optimization of mRNA constructs

Hybridization and opening energies, ΔE_SD_ and ΔE_open_, were calculated with *RNAduplex* and *RNAup*, respectively ^**[15,16]**.^ ΔE_tRNA_ is added as a stabilizing constant, -1.19 kcal/mol or -0.075 kcal/mol only for start codons AUG or GUG, respectively. Optimization is conducted with a self-devised simulated annealing algorithm that performs, selectively accepts and characterizes random single-nucleotide swaps in source transcripts. ΔE_open_(i) for single and sets of constructs, respectively, were calculated with *RNAplfold* from genome sequences available at *microbes.uscs.edu* and *ensemble_biomart* ^**[14,17]**.^

## Results and Discussion

The design of constructs is such that enables not only to understand and assess the influence of expression modulators on protein expression efficiency, but also to assign their optimal location upstream and downstream the initiation codon. The generation of constructs was accomplished by the recombination of a PCR-product into commercially available plasmids. The PCR-product consists of a specific nucleotide sequence or *primer*, and the VDAC-encoding sequence. By the introduction of self-designed primers we are able to modify the genetic code in a controlled fashion and hence assess the effect of these modifications on protein expression. The original pDEST17 plasmids provide sequences before (upstream) and after (downstream) the ATG codon in the untranslated and translated regions, 5’UTR and TR, respectively (figure 1a). The UTR is preceded by the T7 promoter and lodges a prokaryotic RBS in the form of a Shine-Dalgarno (SD) sequence. The TR starts with a 26-amino acid-long sequence containing a 6x HIS-tag (figure 1b). pDEST17 allows the insertion of the PCR-product right after this sequence in the TR. Consequently, the 5’UTR and the location of the RBS is fixed.

**Figure 1.**
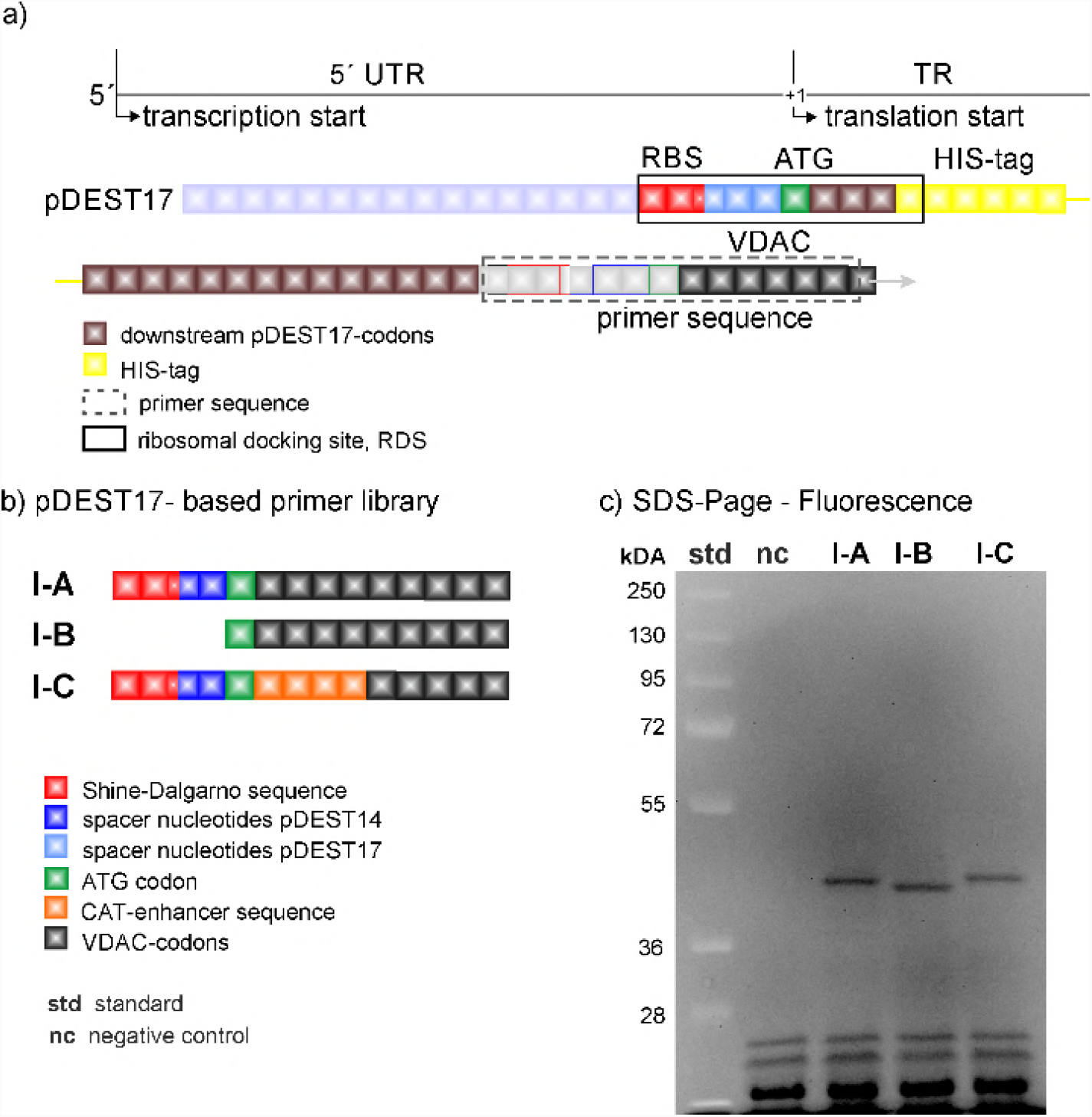
The nucleotide sequence of pDEST17-based plasmids in the proximity of the start codon (a), primer library (b); expression levels (SDS-page gel fluorescence scan) (c); nc: negative (no plasmid) control.

Figure 1c shows that pDEST17-based constructs (VDAC-I) enable protein expression, and alterations in the genetic code far downstream the start codon have no significant effect on protein expression efficiency. The SDS-page gel and Western Blot ^[18]^ of the reaction mixture after protein expression with constructs VDAC-I-A and VDAC-I-B and in the presence of fluorescent lysine, display a single band at approx. 39 kDa of similar intensity. This is indicative of similar expression levels of a single protein. Protein characterization via MALDI-TOF mass spectrometry of trypsin-digested protein fragments, reveal that 34% of the peptides match VDAC sequences, which proves sufficient to confirm the primary structure of VDAC ^[19]^. The insertion of the chloramphenicol acetyltransferase (CAT)-enhancer sequence, ^[20]^ as in VDAC-I-C, does not significantly increase the level of protein expression. The results point to the HIS-tag-containing sequence, possibly in combination with the RBS-starting sequence, as the essential cause for VDAC expression.

Our first hypothesis sets the length and nature of the untranslated region between the T7 promoter and the start codon, the 5’ UTR, as decisive in gene transcription and translation, and we thus employed the plasmid pDEST14 to gain better control over this region, while aiming to express native, tag-free VDAC at comparable levels to those attained through pDEST17-based constructs. pDEST14 provides the T7 promoter as its pDEST17 counterpart does but, unlike the latter, it allows the insertion of self-designed primers at desired locations upstream and downstream the start codon.

Figure 2 shows the primer sequences of the pDEST14-based constructs (VDAC-II) and their respective VDAC expression levels. Though the primer sequences of VDAC-II-A and VDAC-II-B are in turn identical to those of VDAC-I-A and VDAC-I-B, there is hardly evidence of protein expression, as shown by the SDS-page gel fluorescence scan (figure 2c) and the corresponding Western Blot ^[18]^. This evinces the enhancer role of the pre-VDAC sequence and the 5’UTR in pDEST17-based plasmids.

**Figure 2.**
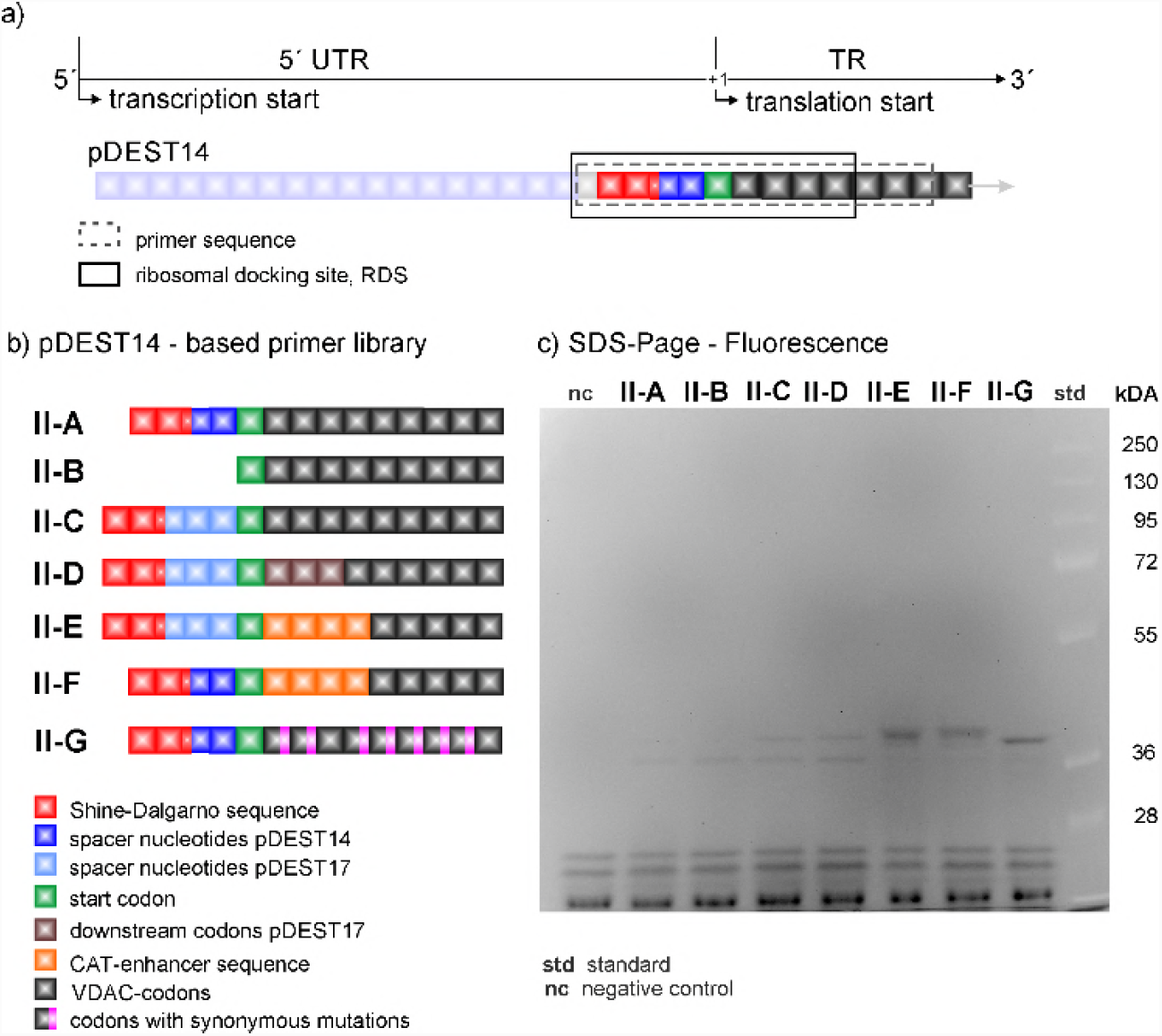
The nucleotide sequence of pDEST14-based plasmids in the proximity of the start codon (a); primer library (b); expression levels (SDS-page gel fluorescence scan) (c) nc: negative (no plasmid) control.

In view of these results, we directed our efforts in investigating the role of the 5’UTR and the adjacent TR in protein expression. Starting at the SD sequence, we inserted the 5’UTR of the pDEST17 vector into pDEST14-based constructs at the same location. The resulting construct, VDAC-II-C, enables marginal protein expression, as evinced by the appearance of a weak band *above* 36 kDa (figure 2c) ^[21]^. So does the construct VDAC-II-D, with the same first three-codon-long coding sequence of the pDEST17 vector. Only the insertion of the 4-codon-long CAT-enhancer sequence after the start codon, as in VDAC-II-E and F, increases the levels of protein expression, irrespectively of the 5’UTR choice. In this case however, VDAC expression is enhanced at the expense of capping the N-terminal of the protein sequence with the non-native amino acid sequence EKKI.

At this point it is crucial to consider whether cell-free VDAC expression is hampered at the transcriptional or translational level. Should gene transcription determine protein expression, mRNA levels would be significantly higher in those cases where protein is expressed, than in those where expression is marginal or not detected. In other words, any changes in levels of transcription by T7 polymerase should result in changes in levels of protein expression. Figure 3 shows that this is not the case; quantitative PCR (Cq values) of cDNA derived from transcripts of different plasmids evince similar levels of mRNA, irrespectively of the plasmid’s translatability, cDNA dilution, and choice of PCR-primer pairs. This and the previous results suggest translation, in particular translation initiation rather than transcription, as the decisive step in determining protein expression, which reverts the focus on the transcript sequence in the immediacy of the start codon.

**Figure 3.**
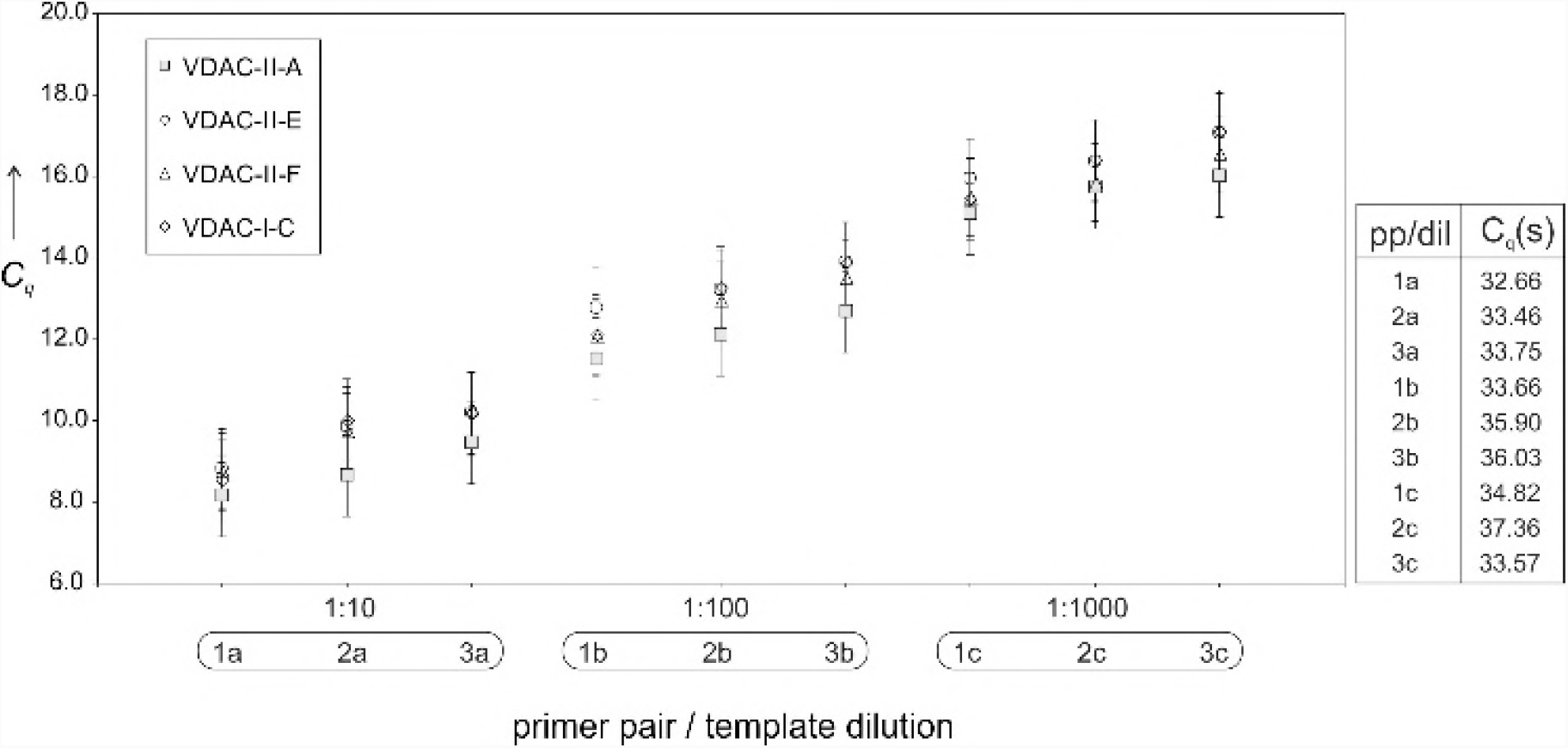
mRNA transcription levels (quantitation cycles, Cq) of different plasmids. Real-time amplification of 3 different dilutions of the as-obtained reverse-transcripted cDNA (a: 1:10; b: 1:100; c: 1:1000) with 3 different primer pairs (1,2, and 3). Right: Cq values for the negative (no plasmid) control.

In contrast to eukaryotic-based expression systems, the prokaryotic machinery is not capable of clearing conformational elements of mRNA that may potentially hamper the correct assembly of the ribosome and hence of the initiation complex ^[22, 23]^. Although the specificity of the interaction between ribosome and mRNA is mediated by hybridization of the SD sequence and strengthened by the coupling of the first transfer-RNA (tRNA^Met^) to the start codon, the whole initiation complex extends over a much longer nucleotide segment. This segment or ribosome docking site (RDS) extends over 30 nucleotides downstream the SD sequence ^[7]^. Since SD sequences are usually positioned 5-13 nucleotides before the start codon ^[22]^, the RDS extends into the coding sequence. Based on this fact, we changed our strategy of ameliorating constructs and opted for a quantitative approach. Inspired by the work of Na *et al*, ^[7]^ we developed a simple *in-silico* translation-initiation potential model to quantify the likelihood of translation of a given mRNA sequence from a series of interaction energy parameters. The model defines the translation-initiation potential δ as

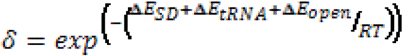

where R is the Boltzmann constant, T the temperature, ΔE_SD_ the hybridization energy between the SD and anti-SD sequences, ΔE_tRNA_ the hybridization energy of the start codon and its respective anti-codon (i.e, the tRNA^Met^), and ΔE_open_ the energy required to unfold the 30-nucleotide-long RDS. Here, ΔE_SD_ and ΔE_tRNA_ are constant since neither the SD nor the start codon are altered. Consequently, variations in δ are exclusively determined by ΔE_open_. Applying the model to the plasmids under study enabled us to rationalize translation events, as translatable constructs consistently scored higher, δ or lower ΔE_open_, than non-translatable ones. Figure 4a shows ΔE_open_ as a function of the position of the SD sequence relative to the start codon, i. The graph displays a minimum at about 11 nucleotides upstream from the start codon only in the case of translatable plasmids (Figure a). The deeper the minimum, the likelier the occurrence of protein expression.

**Figure 4.**
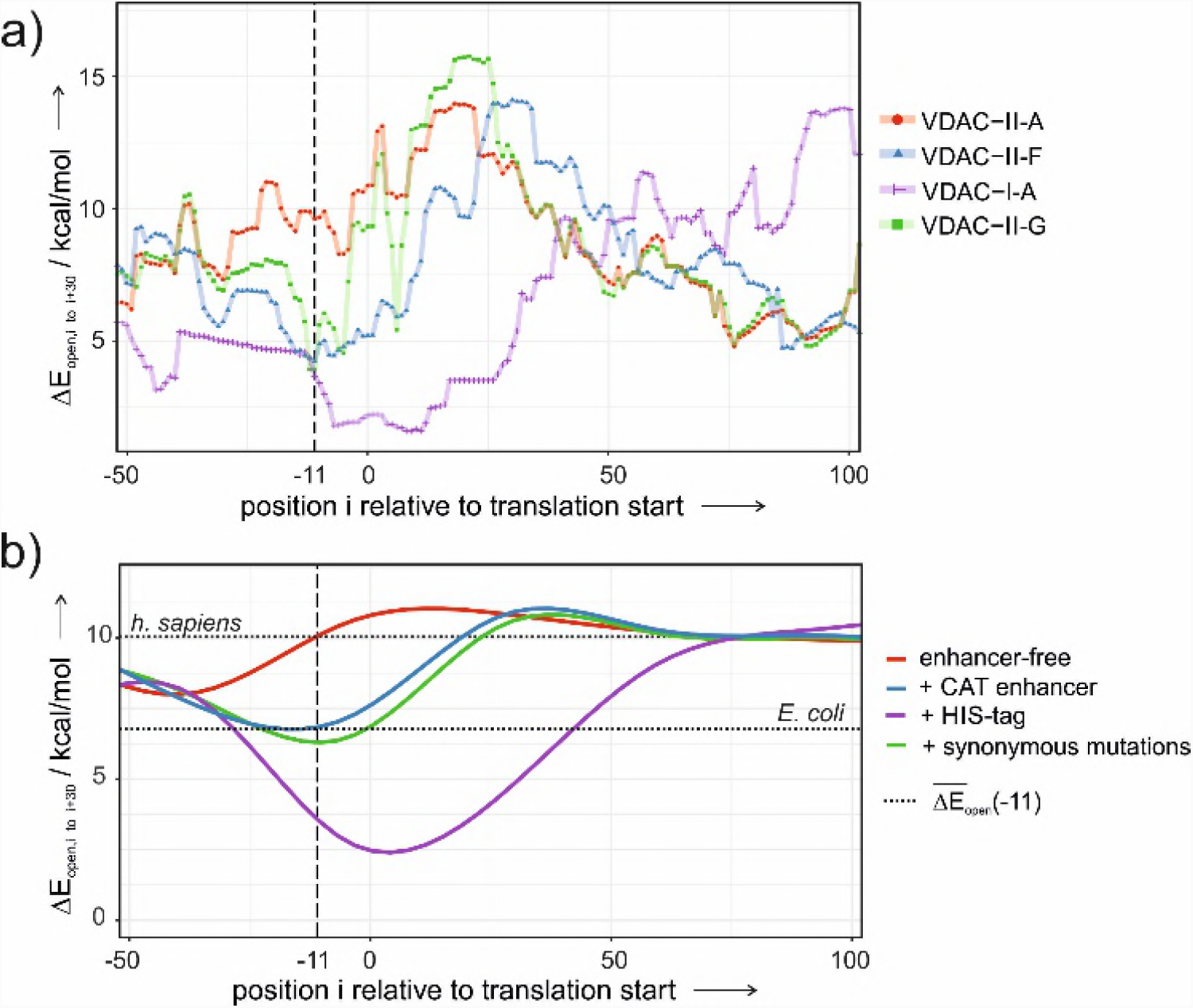
ΔE_open_(i) to unfold the 30-nucleotide-long RDS starting at location i with respect to the start codon (i=0). a) Translatable (VDAC-II-F,G, VDAC-I-A) and non-translatable constructs (VDAC-II-A) for VDAC expression. b) Average ΔE_open_(i) for pDEST14 constructs of all human membrane proteins with and without expression enhancers, and after coding sequence optimization with synonymous mutations. Dotted lines depict the values of ΔE_open_ at i = -11 for the human and *E. coli* genomes.

In view of these results, ΔE_open_ was used as scoring function in a simulated-annealing algorithm to obtain a sequence that maximizes the accessibility of the RDS, preserves the 5’UTR and the native VDAC-coding sequence. Henceforth, we exploited the redundancy of the genetic code by introducing single-nucleotide, synonymous mutations in the VDAC-coding sequence. Applying the optimization algorithm on VDAC-II-A results in the construct VDAC-II-G, sporting seven synonymous mutations in the first nine TR codons. VDAC-II-G encodes the wild-type amino acid sequence of VDAC, and displays a low value of ΔE_open_ at the right location (Figure 4a). Figure 2c shows that VDAC-II-G experimentally enables protein expression in a comparable degree to those attained with enhancer-containing sequences.

Cell-free protein synthesis is governed by the biochemical conditions and the template DNA sequence. The *E. coli*-based system used in this study requires high concentrations of phage T7-RNA polymerase and a surplus of fast degradable amino acids, such as arginine, cysteine, tryptophan, glutamate, aspartate, and methionine ^[24, 25]^. Though necessary, these conditions are not as crucial in protein expression as the mRNA sequence, or rather, the mRNA conformational structure. Sequence elements in the proximity of the start codon, either upstream or downstream, are known to significantly affect translation efficiency ^[26,27,28]^ Which not only implies finding the optimal location for the RBS ^[29]^, but also proper tailoring of the whole RDS. Our findings are based on the design of several plasmids in which the sequence in the immediacy of the start codon have been altered to accommodate the RBS and the gene of a membrane protein at varying distances upstream and downstream the start codon, respectively. The results so far indicate that the best strategy to elicit tag-free protein expression from constructs with off-the-shelf RBSs in prokaryotic cell-free expression systems entails proper engineering of the TR proximal to the initiation codon.

Translation initiation in prokaryotes differs from that of eukaryotes in that it involves much less molecular factors and is significantly less complex. As pointed above, prokaryotes lack mRNA unfolding mechanisms that facilitate the formation of the translation initiation complex, and hence are expected to rely on low-ΔE_open_ transcripts to ensure the expression of their genes.Indeed, ΔE_open_ at i ~ -11 is significantly lower for transcripts of the *E. coli* genome than for those of the human genome ^[30]^. On the other hand, upregulation mechanisms for protein expression in prokaryotic cells are not present in cell-free systems, and may be responsible for in-cell expression of recombinant VDAC from plasmids that do not elicit expression otherwise [31]. Hence, the mRNA sequence is crucial in the cell-free context. Since the ribosome footprint on the mRNA sequence is larger than the RBS and extends well into the TR, a correspondingly long mRNA segment should be accessible for the ribosome to properly dock at and initiate translation. Hence, it makes sense to modify the mRNA sequence within the proximal TR so as to prevent the formation of hindering conformations and gain full access to the RDS. A low ΔE_open_(−11) can thus be viewed as a *sine qua non* criterion for cell-free protein expression with a prokaryotic machinery. According to figure 4a, the efficiency in VDAC expression varies with the nature of the construct as follows: I-A > II-F ≅ II-G >> II-A. A trend that has been qualitatively confirmed by the experiments (figures 1c and 2c).

In this line, the role of translation enhancers in constructs with prokaryotic-like UTRs can be explained. Inserting human genes into pDEST14 vectors alone does not result in values of ΔE_open_ low enough to allow expression (figure 4b, red curve). Contrarily, the insertion of the 6xHIS-tag or the CAT enhancer nucleotide sequences significantly reduces ΔE_open_ to similar or lower values than those of *E. coli* transcripts (figure 4b, blue and purple curves). Enhancers thus enable membrane protein expression inasmuch as they facilitate ribosome assembly through a less structured mRNA in the proximal TR.

Though both enhancers appear as valid options for constructs with poor translation efficiency, they may not be so in those cases where proteins with bare N-terminals and native amino acid sequences are required ^[32]^. Fortunately the redundancy of the genetic code provides enough maneuverability to reduce ΔE_open_ without altering the amino acid sequence, as shown in the case of VDAC. A potentially working strategy that can be applied to any human membrane protein with prohibiting high ΔE_open_ transcripts, by reducing the magnitude to permissive prokaryotic values (figure 4b, green curve). Our results hence suggest that construct optimization via synonymous mutations can be effectively employed in all those other cases where poor mRNA accessibility compromises the outset of translation and hence protein expression.

The current study demonstrates that prokaryotic cell-free expression of VDAC is determined by the mRNA sequence in the immediacy of the start codon and its impact on translation initiation. Providing the RBS site is optimal, i.e., 11 nucleotides upstream the ATG codon, the efficiency of protein expression can be enhanced by introducing synonymous mutations *in the first 9 codons of the TR*. Computer calculations have provided the scoring function ΔE_open_ that allows the quantitative assessment of the translation initiation potential for the plasmids herein investigated, and an optimized, enhancer-free DNA sequence that allows the cell-free expression of native VDAC. This computerized approach can thus predict the performance of plasmids in cell-free protein expression, and provide the optimized sequence of translatable plasmids for VDAC and other human membrane proteins. Though challenges still remain concerning function and structure of membrane proteins, our study smooths the path for an effective go in membrane protein synthesis.

## Acknowledgements

The authors thank Joel Chopineau for kindly providing the plasmid vector pQE30-VDAC. FA thanks the Austrian Science Fund (FWF) for its financial support (Project SFB F43). We are acknowledging the Open Access fond of the University of Natural Resources and Life Sciences, BOKU, Vienna, for financial support.

